# AXL mediates mast cell survival and resistance to tyrosine kinase inhibitors in mastocytosis

**DOI:** 10.1101/2025.11.03.686205

**Authors:** Kitsada Kangboonruang, Philippe Drabent, Fatlinda Maksut, Maxime Heintzé, Yves Lepelletier, Ludovick Lhermitte, Mélanie Feroul, Sébastien Letard, Caroline Kaboré, Fabienne Brenet, Cécile Méni, Nicolas Cagnard, Vincent Bondet, Guillaume Lefèvre, Julie Bruneau, Michael Dussiot, Héloïse Halse, Amélie Bigorgne, Anne-Florence Collange, Hassiba Bouktit, Frédérique Rétornaz, Jérome Megret, Stéphane Barete, Nathalie Droin, Cristina Bulai Livideanu, Angélique Lebouvier, Darragh Duffy, Eric Solary, Michel Arock, Damaj Gandhi, Christine Bodemer, Julien Rossignol, Laura Polivka, Thierry Molina, Olivier Hermine, Leïla Maouche-Chrétien

**Affiliations:** Laboratory of Cellular and Molecular Mechanisms of Hematological Disorders and Therapeutic Implications, Inserm U1163, Institut Imagine, Université Paris Cité, Paris, France; Department of Pathology, Hôpital Necker-Enfants Malades, AP-HP, Université Paris Cité, Paris, France; Laboratory of Genome Dynamics in the Immune System, INSERM 1163, Institut Imagine, Université Paris Cité, Paris, France; Onco-Hematology Department, Hôpital Necker-Enfants Malades, Assistance Publique-Hôpitaux de Paris-AP-HP, Paris, France; Centre de Recherche en Cancérologie de Marseille, INSERM U1068, Marseille, France; Centre de Recherches en Cancérologie de Toulouse (CRCT), INSERM U1037, Team METAML - Metabolism and Drug Resistance In Acute Myeloid Leukemia, Toulouse, France; Department of Dermatology, Hôpital Necker -Enfants Malades, AP-HP, Université Paris Cité, Paris, France; Bioinformatics Department, IMAGINE Institute, Université Paris Descartes, Paris, France; Translational Immunology Unit, Institut Pasteur, Université Paris-Cité, Paris, France; University of Lille, CHU-Lille, Lille, France; CEREMAST, AP–HP, Hôpital Universitaire Necker-Enfants Malades, Université Paris Cité, Paris, France; Service de Médecine Interne et Maladie Infectieuse, Hôpital Européen de Marseille, France; Plateforme de Cytométrie de la SFR Necker, Université Paris Cité, Paris, France; CEREMAST, the Department of Dermatology, Pitié-Salpêtrière Hospital, AP-HP, Paris, France; Université Paris-Saclay, Gustave Roussy, Inserm, Cellules Souches Hématopoïétiques et Développement des Hémopathies Myéloïdes, F-94805, Villejuif, France; CEREMAST, the Department of Dermatology, Hôpital Larrey, CHU Toulouse, Toulouse, France; CEREMAST, the Hematology Institute, Normandy University School of Medicine, Caen, France; Hematology Service, Hôpital Necker-Enfants Malades, AP-HP, Université Paris Cité, Paris, France

**Keywords:** Mastocytosis, mast cell, *KIT*, *AXL*, oncogenesis, cooperation, TKI resistance

## Abstract

Mastocytosis is a clonal disorder driven by *KIT* mutations, but resistance to tyrosine kinase inhibitors (TKIs) remains a major challenge. Following the discovery of an *AXL* L197M mutation in a patient with congenital aggressive mastocytosis, we demonstrated unexpected wild-type AXL expression in neoplastic mast cells (MCs) across mastocytosis subtypes, challenging current views concerning mastocytosis pathophysiology. AXL was undetectable in steady-state MCs but several factors, including IFN-α and IFN-β, induced its expression, consistent with the inflammatory nature of mastocytosis and the high interferon levels in patient plasma.

Ectopic expression of WT or L197M *AXL* in the ROSA *KIT* D816V cell line enhanced proliferation and survival by upregulating pSTAT5, pSTAT3, pFAK, p-p38α, survivin and BCL2. Both AXL forms conferred resistance to the KIT inhibitor PKC412/midostaurin by sustaining BCL2, MCL1, and BCL-XL expression while reducing caspase-3 activation. L197M *AXL* induced slightly stronger resistance to apoptosis than WT, but this difference was not significant. Combined KIT and AXL targeting (PKC412+R428) restored TKI sensitivity by downregulating BCL-XL, Livin and cIAP1, and activating caspase-3, highlighting the therapeutic potential of dual KIT/AXL pathway inhibition. Importantly, neoplastic MCs from a mast cell leukemia patient harboring the *KIT* F522C mutation and unresponsive to PKC412 strongly expressed AXL and displayed marked *in vitro* sensitivity to R428 alone, highlighting AXL as a potential therapeutic target in aggressive mastocytosis not driven by *KIT* D816V.

These findings identify AXL as a previously unrecognized driver of malignant MC survival and TKI resistance, and support AXL inhibition as a promising therapeutic strategy in aggressive mastocytosis.

**Key Points:**

– AXL is aberrantly expressed in neoplastic mast cells, driving survival and resistance to KIT inhibition in mastocytosis.
– Dual KIT and AXL inhibition restores TKI sensitivity in KIT-mutant mastocytosis.

**Graphical abstract:** 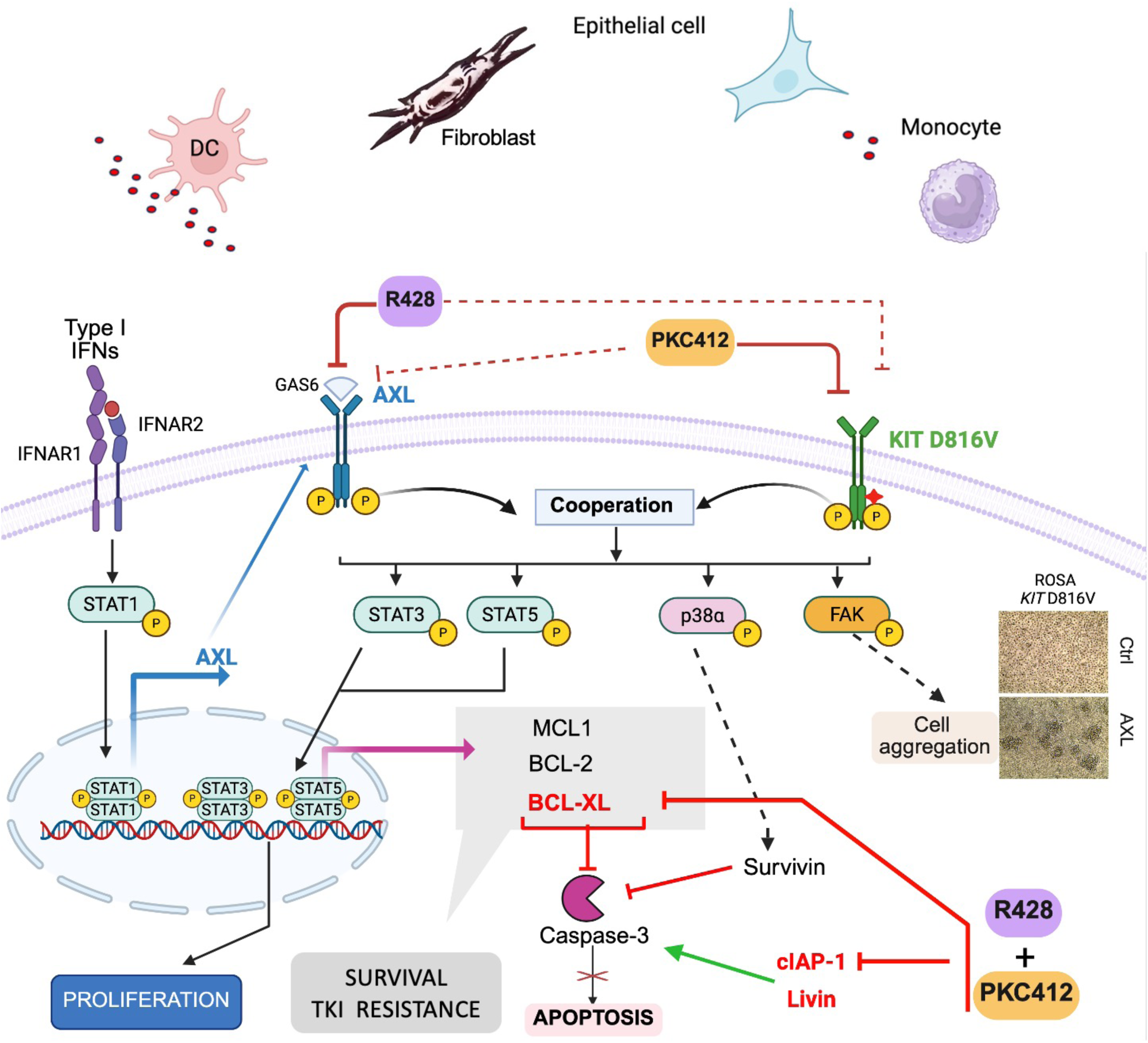

## Introduction

Mastocytosis is defined by multifocal clonal expansion of mast cells (MCs), with KIT receptor tyrosine kinase mutations detected in ∼85% of cases. MC expansion may be confined to the skin (cutaneous mastocytosis, CM), restricted to the bone marrow (BMM), or involve multiple tissues in systemic mastocytosis (SM). SM encompasses indolent (ISM) and smoldering (SSM) forms, both associated with near-normal life expectancy but impaired quality of life due to MC mediator release. Advanced SM (AdvSM) includes aggressive SM (ASM), mast cell leukemia (MCL), and SM with an associated hematologic neoplasm (SM-AHN), all linked to markedly decreases in survival ^1–3^. Additional mutations in *TET2, ASXL1, SRSF2, NRAS,* and *RUNX1* confer poor prognosis in AdvSM ^4–6^. Outside these rare cases, resistance to tyrosine kinase inhibitors (TKIs) is common, and complete responses occur in only 30% of patients, highlighting the need to explore the genetic landscape of mastocytosis beyond *KIT* to shed light on disease heterogeneity and treatment resistance.

We explored familial and congenital forms of mastocytosis and identified a heterozygous germline missense mutation (L197M) in the *AXL* gene of a pediatric patient with congenital ASM. AXL, a member of the TAM receptor tyrosine kinase family, plays crucial roles in hematopoiesis, immune regulation and apoptotic cell clearance ^7^. AXL has been implicated in various cancers, including cutaneous melanoma, squamous cell carcinoma, CLL, and AML, in which it is recognized as a key mediator of TKI resistance. In most tumors, AXL exerts its oncogenic functions through expression or overexpression of the wild-type receptor, while mutations are rarely reported ^8–10^. Under physiological conditions, AXL is expressed in multiple cell types, particularly myeloid cells, such as monocytes, macrophages, and dendritic cells ^11,12^. Its canonical activation is mediated by the ligand Gas6, although alternative mechanisms have also been described ^13^. Ligand-dependent or -independent activation induces AXL autophosphorylation, triggering downstream signaling pathways promoting proliferation, immune evasion, apoptosis resistance and EMT ^14,15^.

Given the well-established role of AXL in oncogenesis and TKI resistance, and the absence of studies on MCs and mastocytosis, we investigated the potential involvement of the wild-type receptor and the germline L197M variant in mastocytosis pathophysiology.

## Materials and Methods

See Supplementary Materials for full methods

### Patient samples

This retrospective study was conducted within the CEREMAST network on BM, skin, and blood samples from mastocytosis patients. It was approved by the local ethics committee (Comité de Protection des Personnes Ile-de-France, approval 93-00) and conducted in accordance with the Declaration of Helsinki; all patients gave written informed consent. Patient data, diagnoses, and *KIT/AXL* status are provided in Table 1

**TABLE 1:**
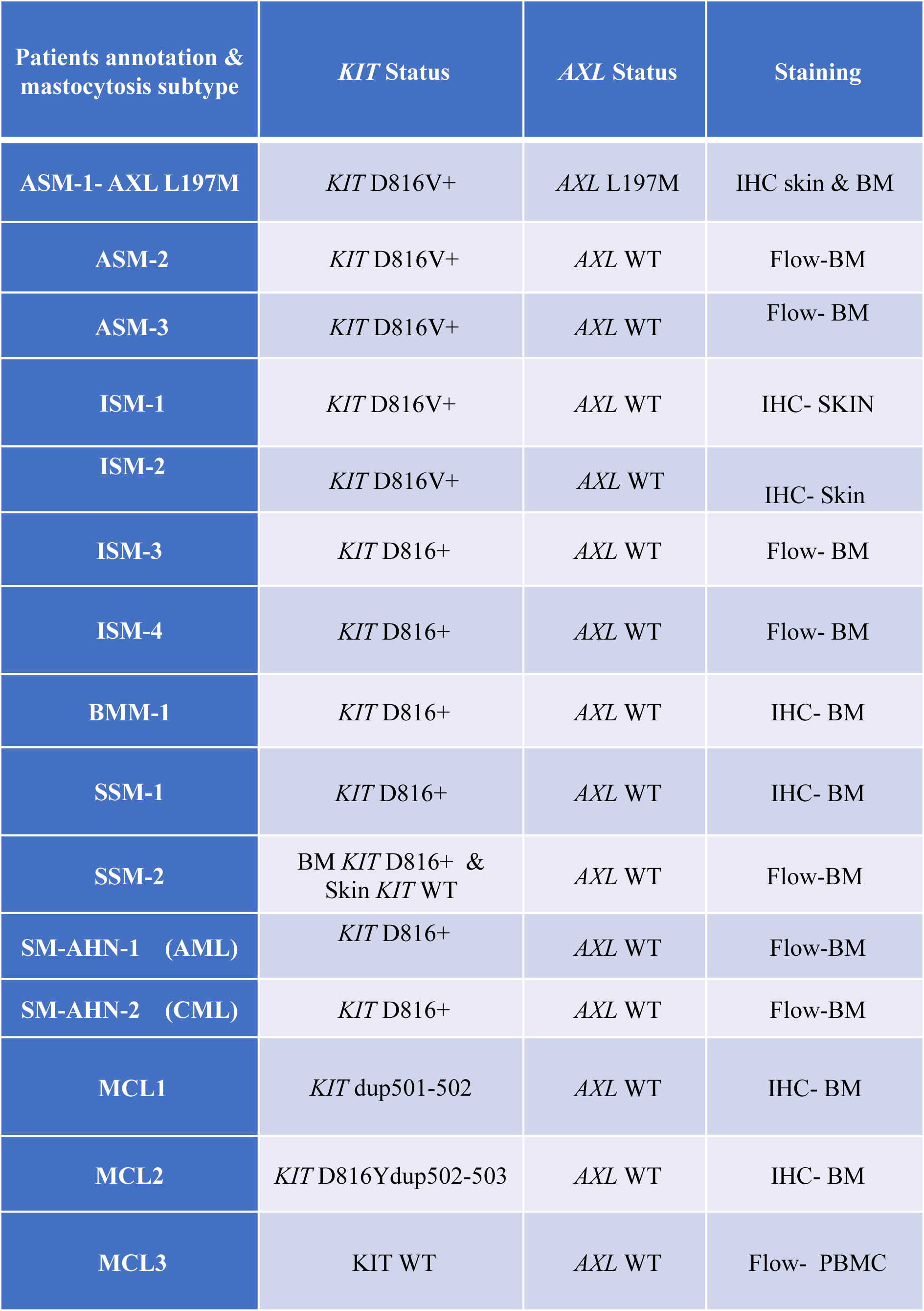
Patient characteristics including mastocytosis subtype, KIT and AXL mutational status, and mast cell detection method (FACS or IHC).

### Immunohistochemical analysis

Skin and BM sections were stained with H&E or subjected to IHC for CD117 and AXL.

### Flow cytometry analysis of AXL expression in MCs

BM samples from mastocytosis patients were thawed, fixed, permeabilized and stained with anti-AXL, anti-CD117, and anti-FcεRI.

### Simoa digital ELISA

Plasma samples from mastocytosis patients were analyzed for IFN-α, -β, and -γ levels by Simoa digital ELISA (Quanterix) and compared with healthy control plasma.

### Plasma GAS6 determination

GAS6 was determined by Luminex assays in 37 SM patients and 10 healthy donors.

### RNA extraction and RT-qPCR

Total RNA was extracted from MC lines and CD34⁺-derived MCs (± IFN stimulation), reverse-transcribed, and analyzed by TaqMan real-time PCR. Expression was normalized against 18S rRNA.

### Mast cell lines

ROSA *KIT* WT SCF-dependent, ROSA *KIT* D816V (CD34⁺–derived) ^16^, HMC1.2 (*KIT* V560G/D816V, MCL patient), and BMMCs from *KIT* D814V transgenic mice were used.

### Primary MCs

MCs were generated from peripheral blood CD34⁺ cells cultured with SCF and IL-6. MC differentiation was confirmed by CD117 and FcεRI expression.

### Lentiviral vector production and cell transduction

The lentiviral vectors pRRSLIN-*AXL* WT-tdTomato and pRRSLIN-*AXL* L197M-tdTomato (backbone pRRSLIN-MND-PGK-tdTomato-WPRE, gift from Prof. F. Moreau-Gaudry) were generated and used to transduce ROSA *KIT* WT, ROSA *KIT* D816V and BMMC *KIT* D814V cells. Two independent batches of transduced cells were analyzed.

### Cell proliferation

The proliferation kinetics of MC lines (ROSA *KIT* WT, ROSA *KIT* D816V and BMMC *KIT* D814V) expressing *AXL* WT-tdTomato, *AXL* L197M-tdTomato or control were monitored for 7 days by IncuCyte, as described elsewhere ^17^.

### RNA-seq on microdissected cutaneous MCs

Cutaneous MCs were microdissected from 12 ISM and 3 AdvSM patients, and RNA was extracted for sequencing.

### *In vitro* cell proliferation with TKI treatment

The proliferation of ROSA *KIT* WT and ROSA *KIT* D816V cells expressing AXL *WT*, *AXL* L197M, or control was monitored for 7 days by IncuCyte after treatment with DMSO (vehicle) or 200 nM PKC412.

### Colony formation assay

ROSA *KIT* D816V cells expressing *AXL* WT, *AXL* L197M, or control were plated in soft agar, and colonies were quantified after three weeks with ImageJ.

### BaF3 cells viability

IL-3-dependent BaF3 cells were transduced with *AXL* WT, *AXL* L197M or control vector and cultured with or without IL3. Cell viability was assessed after 48 h in Alamar Blue assays (n=3).

### Cell death analysis

ROSA *KIT* D816V cells expressing *AXL* WT, *AXL* L197M, or control were treated with 200 nM PKC412 or DMSO, and cell death was assessed by FACS with SYTOX® Blue staining.

### Transcriptomic analyses

RNA-seq was performed on RNA from ROSA *KIT* D816V cells and from cells expressing *AXL* WT or L197M. Functional analyses were performed by IPA (http://www.ingenuity.com) on the genes identified. The data are available from https://www.ebi.ac.uk/biostudies/studies/S-BSST2177.

### Immunoblotting

Protein extracts from ROSA and HMC1.2 cells ± IFN were probed with anti-AXL. STAT3/5, FAK, p38, BCL-XL, BCL2, MCL1, caspase-3, survivin, cIAP1 and Livin were analyzed in ROSA *KIT* D816V expressing *AXL* WT or L197M and control cells. Cells were left untreated or treated for 48H with DMSO, or 200 nM PKC412 ± 500 nM R428. Blots were normalized against β-actin or HSP70.

### Cell viability assay

ROSA *KIT* D816V cells (control, *AXL* WT, or L197M) were treated for 7 days with DMSO, 200 nM PKC412, 500nM R428, or both. Viability was assessed by WST-8 at 450 nm.

### TKI treatment of MCL3 primary MCs

Fresh PBMCs from the MCL3 patient with *KIT* F522C, *SF3B1* and *TERT* mutations were treated for 3 weeks with DMSO (control), 200 nM PKC412, 500 nM R428, or both. MC survival was assessed by FACS (CD117, FcεRI, Sytox). Small cell numbers limited duplication.

## Results

We identified a pediatric patient with congenital ASM carrying a germline *AXL* L197M missense mutation and a somatic *KIT* D816V mutation. PolyPhen and SIFT scores of 0.986 and 0, respectively, suggested a potentially damaging effects on protein function **(Figure 1A)**. However, the CTG-to-ATG variant has never been associated with tumors. Given that oncogenic roles of *AXL* are typically driven by expression of the wild-type form, we investigated the involvement of WT *AXL* and L197M *AXL* in mastocytosis.

**Fig 1.**
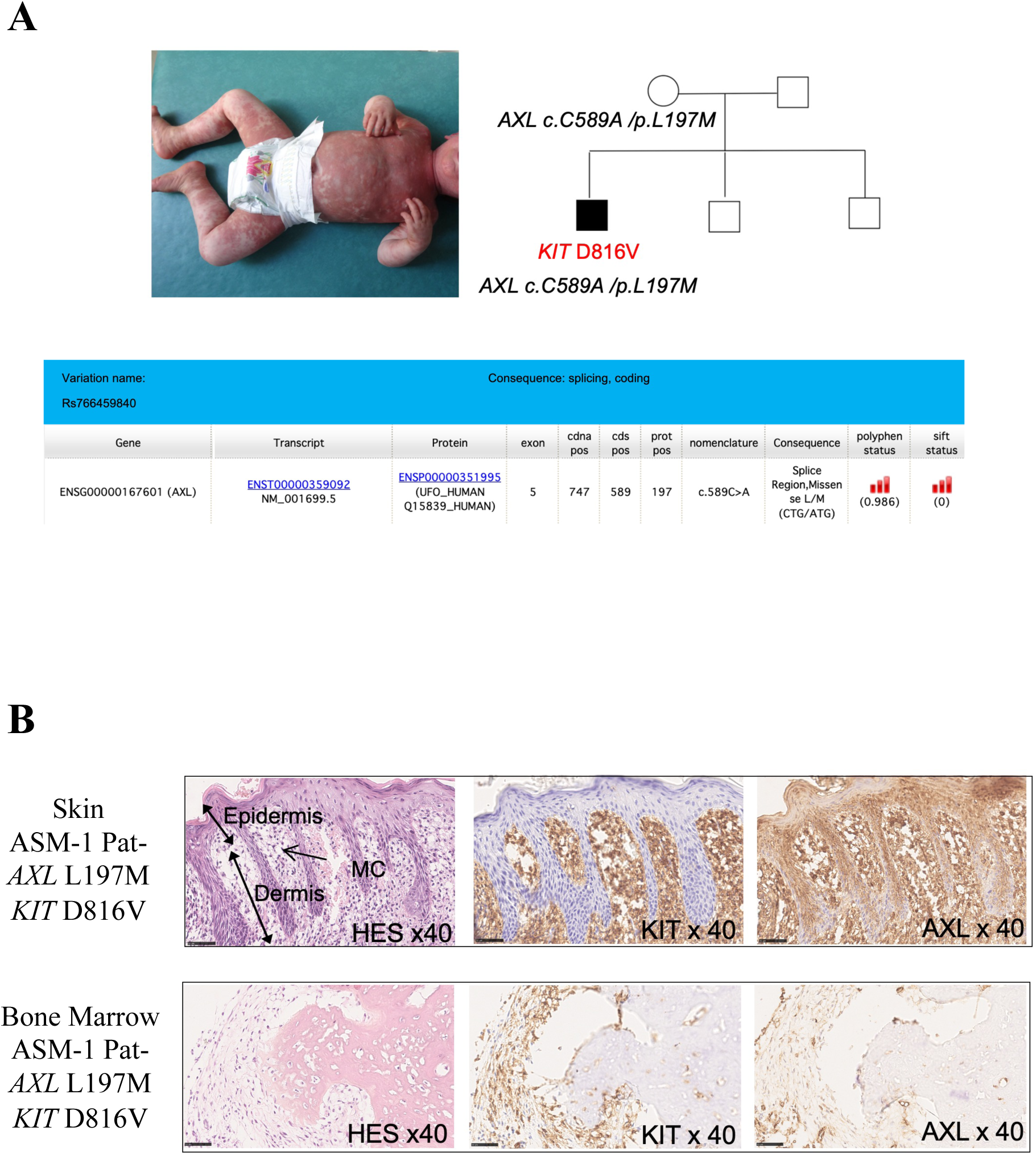
Genetic profile and AXL expression in a patient with congenital aggressive systemic mastocytosis (ASM). **(A)** The patient carries a somatic *KIT* D816V mutation and a germline heterozygous *AXL* L197M mutation. (**B)** Histological analysis of skin and bone marrow (BM) sections highlight KIT positive MC infiltrates. Strong expression of AXL in MCs from skin and moderate in the BM. Original magnification, ×40 (insets).

We first assessed *AXL* expression in MCs by Immunohistochemistry (IHC) on skin and bone marrow (BM) sections from the *AXL* L197M patient (Pat-*AXL* L197M) and from several patients with different degrees of MC infiltration **(Table 1)**. None of the other patients carried *AXL* mutations. Hematoxylin/eosin (H&E) and KIT staining were used to identify MC infiltrates. Pat-*AXL* L197M sections displayed extensive MC infiltration with strong AXL expression, more pronounced in skin **(Figure 1B)**. Marked expression was also observed in one mastocytoma and one ISM skin section analyzed in parallel **(Supplementary Figure 1)**.

Skin sections from two ISM patients (ISM-1 and ISM-2) showed moderate and weak MC infiltration, respectively, with detectable AXL in both **(Figure 2Aa)**. BM sections from patients with various forms of mastocytosis revealed heterogeneous AXL expression. Expression was strong in neoplastic MCs from one SSM and one MCL case (SSM1-MCL-1), weak in a BMM case, and intermediate in a second MCL case (MCL-2) **(Figure 2 Ab)**. Given that AXL is also expressed by myeloid cells, we performed flow cytometry on frozen BM samples from eight patients (2 ISM, 2 SM-AHN, 1 SSM, 2ASM) and fresh blood from one MCL patient to confirm AXL expression in neoplastic MCs. In healthy individuals, bone marrow MCs (CD117⁺/FcεRI⁺) account for <10⁻⁵ of total cells. In our cohort, MC infiltration ranged from 0.02% to 27%, with all MCs expressing AXL, albeit at different levels **(Figure 2B)**. Specifically, expression was low in MCs from ISM-4 (0.3%) and SM-AHN-1 (5%), whereas it was high in MCs from ISM-3 (1.2%), SM-AHN-2 (0.02%), SSM-2 (3%), ASM-2 (0,3%), ASM-3 (2.66%) and MCL3 (27%) **(Figure 2B)**. This variability is consistent with the heterogeneous IHC staining of BM biopsy specimens, although different patients were analyzed with the two methods. RNA-seq analysis of skin lesions from 15 mastocytosis patients (12 ISM, 3 AdvSM) revealed *AXL* expression in all cases, with higher levels in neoplastic MCs from the small AdvSM subgroup **(Figure 2C)**. Given the differential AXL expression in mastocytosis, we assessed plasma GAS6 levels in Luminex assays. No significant difference was found between patients (n=37) and healthy controls (n=10) **(Figure 2D)** ^13^.

**Fig 2.**
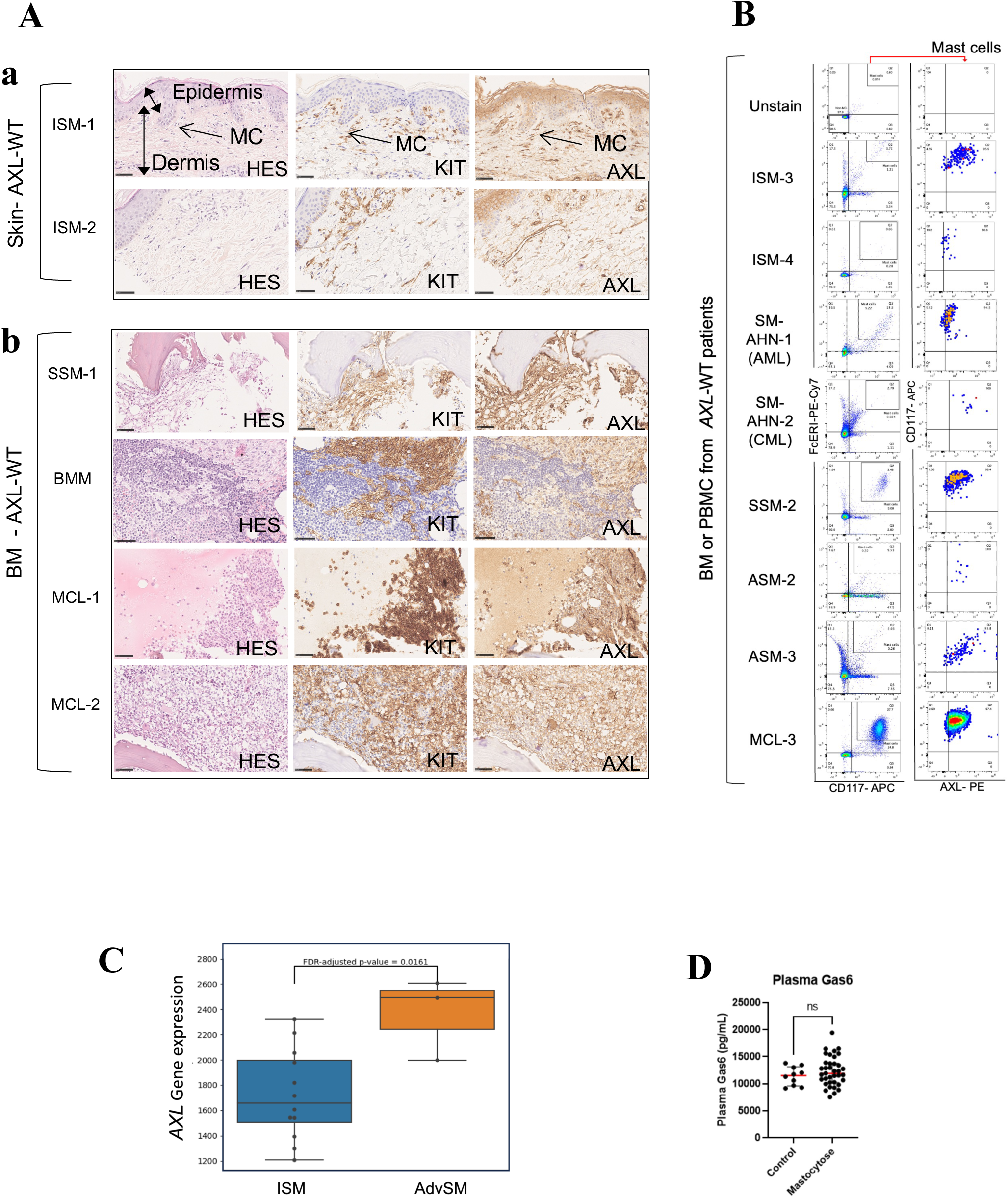
AXL expression in neoplastic MCs across mastocytosis subtypes. **(A)** histological analysis of skin and BM sections from patients with mastocytosis, all wild type for AXL (*AXL*-WT). KIT staining identifies MC infiltrates**. a)** Skin sections from 2 ISM cases show moderate MC infiltrates, all positive for AXL. **b)** BM sections from 1 SSM, 1 BMM and 2 MCL cases show abundant MC infiltrates with variable AXL expression levels. Myeloid cells from MCL2, are KIT negative and AXL positive. Original magnification, x40 (insets). ***(B)*** Flow cytometry staining with anti-(CD117, FceRI and AXL) on 7 DMSO-frozen BM samples from patients: 2 ISM, 2 SM-AHN (AML/CML), 1 SSM, 2ASM and 1 PBMCs MCL sample. ***(C)*** AXL expression assessed in neoplastic MCs sorted from skin lesions of 15 mastocytosis patients (12 ISM and 3 AdvSM), and analyzed by RNA sequencing (p-value=0.0161). ***(D)*** Plasma levels of GAS6 ligand in healthy individuals (n=10) and patients with systemic mastocytosis (n=37), measured using a Luminex assay. (ns: not significant).

### AXL is barely detectable in normal MCs

Assessing *AXL* expression in MCs from normal skin or BM is challenging due to their low abundance and widespread *AXL* expression in non-MC populations. scRNA-seq provides some insight, but fresh biopsies from healthy individuals are difficult to obtain for ethical reasons. We therefore analyzed publicly available scRNA-seq datasets, focusing on the Immune Single-Cell Consortium dataset (*disco_mast_basophil_v1.0.rds*). We identified 2,400 MCs in normal skin from 26 individuals. *AXL* expression was found in only 21 cells (∼0.9%) from three individuals who also had *AXL*-negative MCs. In primary skin tumors from 11 patients (basal and cutaneous squamous cell carcinoma), *AXL* was detected in only 1 of 536 MCs (∼0.2%). In 68 normal BM samples, 1,086 MCs were identified, 74 (∼6%) of which expressed *AXL*. These cells were obtained from six individuals. Overall, *AXL* expression is rare to absent in MCs from normal skin and BM but widely expressed in neoplastic MCs from mastocytosis patients. Given the limited accessibility of MCs from both healthy and diseased tissues, technical hurdles remain for in-depth studies of AXL function in primary MCs. We addressed this issue using ROSA *KIT* WT, ROSA *KIT* D816V, and HMC1.2 mast cell lines carrying wild-type *AXL*.

### Inflammatory factors induce *AXL* expression in mast cells

*AXL* expression was assessed by RT-qPCR in human MC lines. mRNA was barely detectable in ROSA *KIT* WT and ROSA *KIT* D816V cells (Cq = 38.02 ± 0.19 and 34.50 ± 0.68, respectively), and was present at low levels in HMC1.2 cells (Cq = 33.38 ± 0.39) **(Figure 3A)**. We therefore investigated whether environmental factors, including hypoxia, IFNs, TNF-α, GM-CSF, LPS, and poly(I:C) could induce *AXL* expression in MCs, as reported in other cell types ^12,18–22^. Exposure of MCs to hypoxia (1% or 3% O₂), poly(I:C), or GM-CSF did not increase *AXL* expression in MCs (data not shown). By contrast, IFN-α (100 ng/mL) and IFN-β (5 ng/mL) significantly upregulated *AXL*, by 135-fold and 390-fold, respectively, in ROSA *KIT* WT cells and by 30-fold and 45-fold in ROSA *KIT* D816V cells. Increases of 80-fold and 74-fold were observed in HMC1.2 cells **(Figure 3A)**. Western blotting confirmed AXL expression after IFN-α or β stimulation **(Figure 3B)**. IFN-γ (10–400 ng/mL) moderately induced *AXL*, with increases of up to 5-fold in ROSA *KIT* D816V and 10-fold in HMC1.2 cells. LPS (100 ng/mL) induced a 15-fold increase **(Figure 3C)**, whereas TNF-α (150 ng/mL) led to a modest 2-fold increase (**data not shown).** IFN-α, IFN-β, and IFN-γ induced the strongest upregulation of *AXL*, confirmed at the protein level by immunoblotting **(Figure 3D)**. We assessed *AXL* expression in primary MCs derived from CD34⁺ progenitors. *AXL* mRNA was almost undetectable (Cq = 38) in basal conditions but IFN-α (100–300 ng/mL) induced increases of up to 300-fold **(Figure 3E)**, confirmed by flow cytometry **(Figure 3F)**. IFN-α induced STAT1 expression and pSTAT1 (Y701) activation in all MC lines, together with AXL upregulation **(Figure 3G-H)**. These findings are consistent with reports that IFN and LPS induce AXL expression via STAT1 activation ^12, 23^.

**Fig 3:**
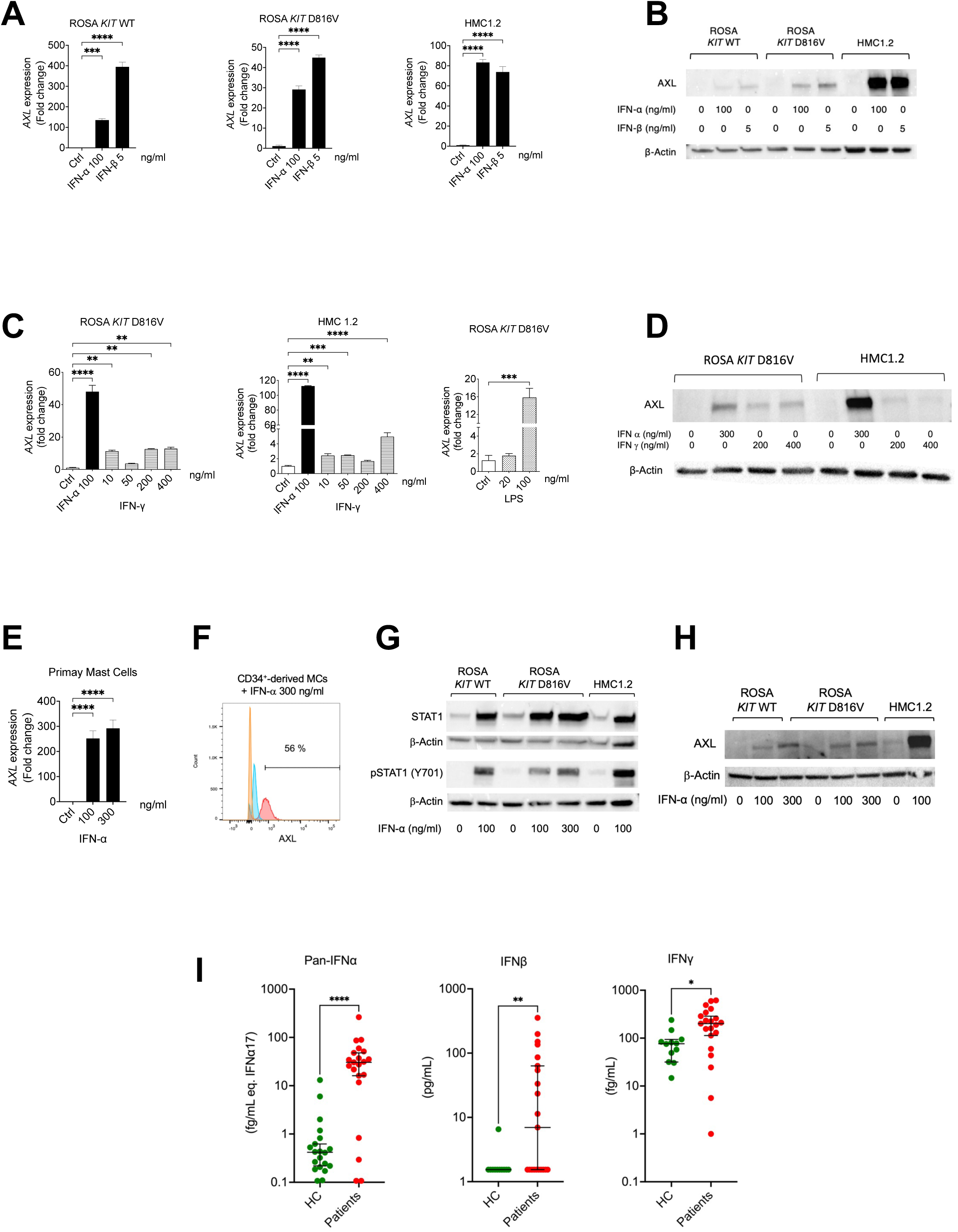
Effect of inflammatory cytokines on AXL expression in mast cells. ***(A)*** RT-qPCR analysis of AXL expression using a specific TaqMan probe in cell lines (ROSA *KIT* WT, ROSA *KIT* D816V, HMC1.2, unstimulated (Ctrl) or stimulated with IFN-α (100 ng/mL) and IFN-β (5 ng/mL). ***(B)*** Immunoblot of AXL in unstimulated MC lines (0) or stimulated with IFN-α (100 ng/mL), IFN-β (5 ng/mL). ***(C)*** AXL expression analyzed by RT-qPCR following IFN-α (100 ng/mL), IFN-γ (10, 50, 200, 400 ng/mL) stimulation in Rosa *KIT* D816V and HMC cells, and LPS stimulation (20, 100 ng/mL) in Rosa *KIT* D816V cells. ***(D)*** Immunoblot of AXL in unstimulated MC lines (0) or stimulated with IFN-α (300 ng/mL), IFN-γ (200–400 ng/mL). ***(E)*** RT-qPCR analysis of AXL expression in primary MCs, derived from CD34+ cells, unstimulated (Ctrl) or stimulated with IFN-α (100–300 ng/mL). All stimulations were performed for 48 hours. RT-qPCR were normalized to 18S rRNA. One-way ANOVA was used for statistical analyses (**** p<0.0001, *\*\*\* =*p<0.001, ** p<0.01*. **(F)*** Detection of AXL in unstimulated or stimulated primary MCs by Flow cytometry (Orange =unstained, Blue= IgG, Pink= AXL). ***(G)*** Immunoblot detection of STAT1 and pSTAT1 (Y701) in MC lines, either unstimulated or stimulated with IFN-α (100 or 300 ng/mL). ***(H)*** Immunoblot of AXL in unstimulated MC lines (0) or stimulated with IFN-α (100 or 300 ng/mL). ***(I)*** Simoa digital ELISA analysis of plasma interferon levels in mastocytosis patients (n=21) and healthy controls: Pan-IFN-α, IFN-β, and IFN-γ. Mann-Whitney test used: **** p<0.0001, ** p<0.01, * p<0.05.

We investigated interferon levels in mastocytosis patients, to assess their involvement in AXL expression in MCs. SIMOA analysis of plasma from 21 patients showed that IFN-α, IFN-β, and IFN-γ levels were significantly higher than in controls **(Figure 3I)**.

### AXL (WT or L197M) enhances *KIT* D816V-driven oncogenicity

MC lines do not express AXL without stimulation. We therefore generated lentiviral vectors encoding *AXL* WT-tdTomato or *AXL* L197M-tdTomato and used them to transduce ROSA cells. Neither WT nor L197M *AXL* expression affected ROSA *KIT* WT cell proliferation, whereas both significantly increased ROSA *KIT* D816V proliferation relative to controls **(Figure 4A)**. Similar results were obtained in BMMC-*KIT* D814V cells, further confirming that both WT and L197M *AXL* enhance proliferation **(Figure 4B)**. AXL expression was verified by immunoblotting in transduced cells **(Figure 4C)**. We then assessed their impact on colony formation in ROSA *KIT* D816V cells. Both WT and L197M *AXL* increased colony numbers, 2.5-fold and 2-fold, respectively, relative to controls **(Figure 4D)**. These findings support oncogenic cooperation between *KIT* D816V and both WT and L197M *AXL*. We investigate the underlying mechanisms by performing immunoblotting for key signaling molecules. Increased pSTAT5, pFAK and p-p38 levels were detected, with a trend toward higher levels of pSTAT3. Total STAT5, STAT3, FAK and p38 were not significantly different after normalization to actin, suggesting that AXL expression is associated with enhanced phosphorylation and not with increased protein abundance **(Figure 4E)**.

**Fig. 4:**
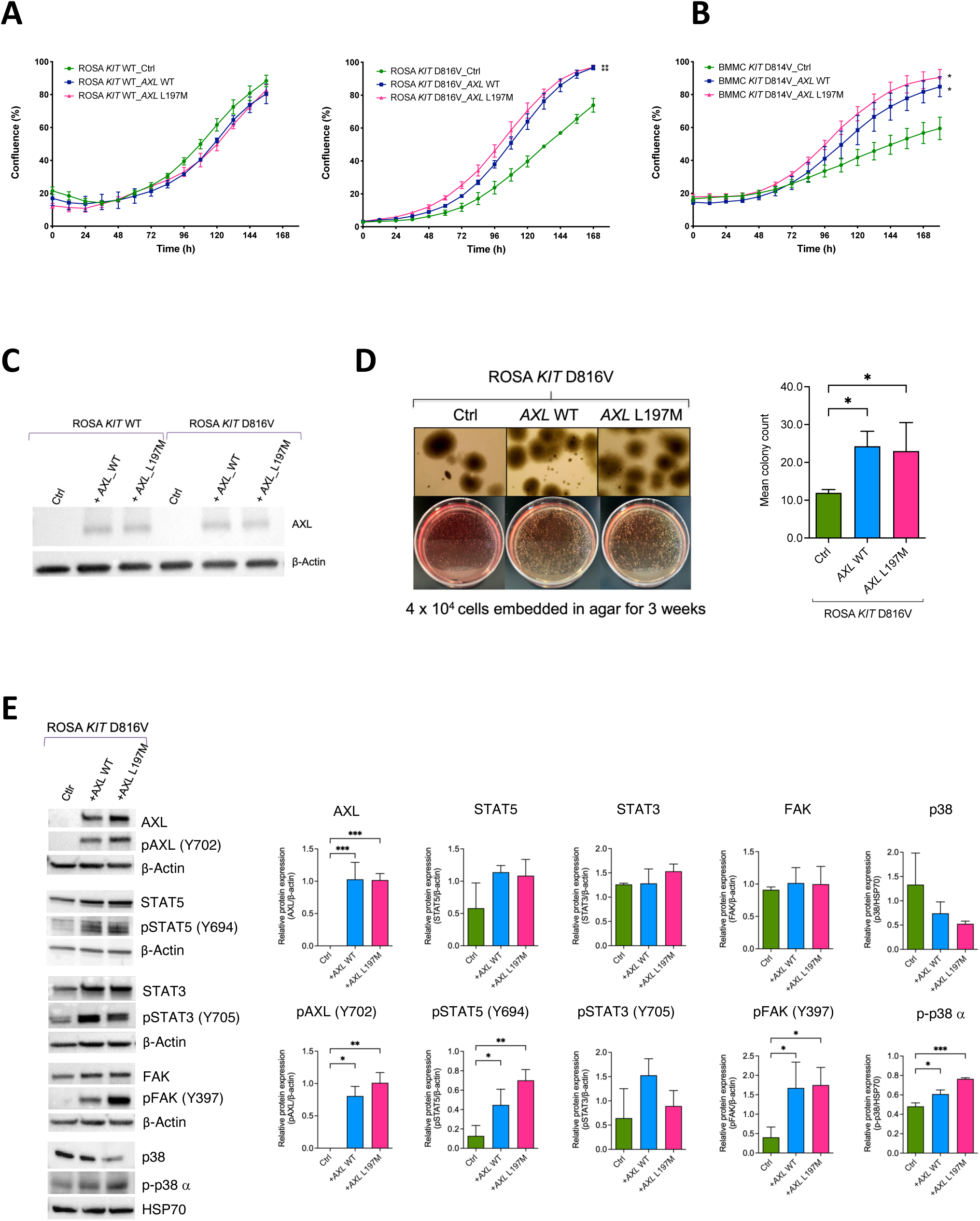
Functional cooperation between D816V KIT and AXL during proliferation. ***(A)*** Proliferation kinetics of ROSA *KIT* WT, ROSA *KIT* D816V, and ***(B)*** BMMC *KIT* D814V transduced with lentiviral vectors *AXL* WT (Blue), *AXL* L197M (Pink) and Ctrl cells (green), monitored by incucyte live-cell imaging. Confluence (%) is plotted against time (in hours). ***(C)*** Immunoblot of AXL in ROSA *KIT* WT and ROSA *KIT* D816V cells transduced with *AXL* WT*-tdTomato*, *AXL L197M*-*tdTomato* lentiviral vectors, or control cells. ***(D)*** Colony formation was assayed in ROSA *KIT* D816V cells transduced with the same lentiviral vectors and control cells. ***(E)*** Immunoblots of AXL, total and phosphorylated STAT5, STAT3, FAK and p38α in transduced ROSA *KIT-*D816V. β-actin or HSP70 were used for normalization. Statistical analyses were performed by one-way ANOVA: *\**p<0.05; **p<0.01; ***p<0.001. Data are representative of 3 independent experiments. é

### AXL confers a survival advantage and induces TKI resistance in *KIT* D816V MCs

ROSA *KIT* D816V cells expressing *AXL* (WT or L197M) consistently showed reduced basal cell death compared with parental cells (∼2% *vs*. ∼5%), indicating a survival advantage **(Figure 5A)**. Accordingly, AXL expression markedly upregulated the survival factors ^24–27^ survivin and BCL2 **(Figure 5B)**. We investigated this survival advantage further in the IL-3-dependent BaF3 cell line, which lacks endogenous *AXL* and *KIT* expression. Neither WT nor L197M *AXL* affected viability in the presence of IL3, but in its absence both conferred a pro-survival effect, with L197M *AXL* having a highest effect (**Figure 5C)**. Neither form of AXL increased the proliferation of BaF3 cells (data not shown), contrasting with the proliferative effect observed in ROSA *KIT* D816V cells **(Figure 4A)**. Transcriptomic analysis comparing parental ROSA *KIT* D816V cells with those expressing WT or L197M *AXL* revealed that expression of either form of XL was associated with an enrichment in pathways related to cancer, metastasis, tumor cell proliferation, cell viability, and survival. In contrast, pathways linked to apoptosis, necrosis, and cell death were downregulated in AXL-expressing cells, consistent with our functional findings **(Figure 5D)**.

**Fig. 5:**
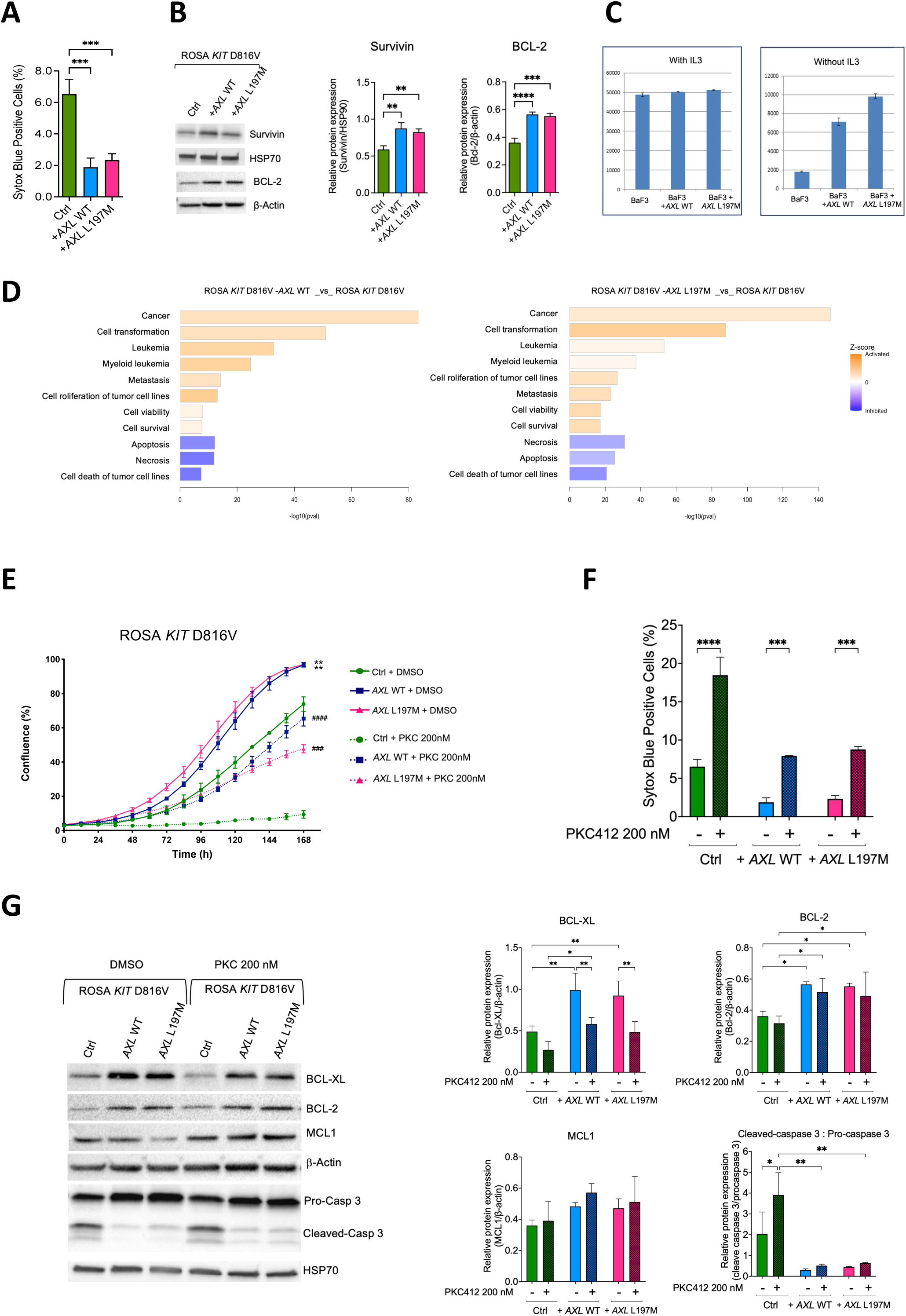
Effect of WT and L197M *AXL* expression on cell survival and PKC 412 treatment. ***(A)*** Proportion of cell death in parental ROSA *KIT* D816V cells and in cells expressing *AXL* WT or *AXL* L197M under basal conditions. ***(B)*** Immunoblot analysis of Survivin and BCL2 in ROSA *KIT* D816V cells expressing AXL WT, AXL L197M, or lacking AXL. Statistical significance was assessed by One-way ANOVA **p<0.01, ***p<0.001, ****p<0.0001***. (C)*** Growth of IL-3-dependent Ba/F3 cells expressing *AXL* WT or *AXL* L197M in presence or absence of IL-3. ***(D)*** Ingenuity pathway analysis (IPA) of RNAseq data from parental ROSA *KIT* D816V cells and counterparts expressing *AXL* WT or *AXL* L197M revealed clustered cellular functions. Activation z-scores were calculated by the IPA z-score algorithm, which predicts the direction of change for a function. An absolute z-score ≥ 2 is considered significant: functions are increased (pink) if z-score ≥ +2 and decreased (blue) if z-score ≤ –2. ***(E)*** Proliferation kinetics of ROSA *KIT* D816V cells expressing *AXL* WT, *AXL* L197M compared with control cells over 7 days of treatment with 200 nM PKC412 or DMSO. One-way *ANOVA* was used: *\*\**p<0.01*: AXL*-expressing cells compared to control treated with DMSO*; ^##^*p<0.01; ^###^p<0.001, ^####^ p<0.0001*: AXL*-expressing cells compared to control cells treated with PKC412*). **(F)*** Cell death measured by SYTOX blue staining after 96-hour treatment with 200 nM PKC412 or DMSO. Significance was determined relative to controls and DMSO by two-way ANOVA (***p<0.001, ****p<0.0001*). **(G)*** Immunoblot analysis of BCL-XL, BCL2, MCL1, and cleaved caspase-3/procaspase-3 ratio in ROSA *KIT* D816V cells expressing AXL WT, AXL L197M, or control, following 48-hour treatment with 200 nM PKC412 or DMSO. Statistical significance was assessed versus DMSO-treated cells using two-way ANOVA (*p<0.05; **p<0.01). Data are representative of ≥3 independent experiments, except cleaved caspase-3/procaspase-3 (n=2).

AXL has been implicated in resistance to TKIs in several cancers, including CML and AML ^8–10^. As up to 70% of ASM patients are resistant to midostaurin (PKC412) and neoplastic MCs express AXL, we hypothesized that AXL might contribute to TKI resistance. Treatment of ROSA *KIT* D816V cells with 200 nM PKC412 inhibited growth by ∼90%, whereas cells expressing WT or L197M *AXL* were less sensitive, with only ∼40% and ∼55% decreases in confluence, respectively (**Figure 5E**). We next evaluated cell death by SYTOX blue staining. Under control conditions (DMSO), *AXL*-expressing cells showed fewer dead cells than parental cells (∼2% vs. ∼6%). After 96 hours of PKC412 treatment, cell death increased in all groups but remained lower in *AXL*-expressing cells (∼7% vs. ∼18%) **(Figure 5F)**. Thus, AXL expression, regardless of the variant, enables *KIT* D816V MCs to evade PKC412-induced growth inhibition and apoptosis, with the L197M variant conferring resistance comparable to WT *AXL*.

We investigated the mechanisms by which WT and L197M *AXL* enhance survival and PKC412 resistance in ROSA *KIT* D816V cells by immunoblotting anti-apoptotic proteins. Both AXL forms were associated with higher BCL-XL, BCL2, and MCL1 levels than control cells, consistent with a prosurvival phenotype **(Figure 5G)**. Treatment with 200 nM PKC412 significantly decreased BCL-XL in all groups, but levels remained high in *AXL*-expressing cells, comparable to untreated controls, indicating relative resistance to PKC412-induced downregulation. By contrast, PKC412 did not affect BCL2 or MCL1, irrespective of AXL expression **(Figure 5G)**. We then assessed caspase-3 activation. In untreated cells, the cleaved/pro-caspase-3 ratio was ∼5-fold higher in control cells than in WT- or L197M *AXL*-expressing cells. PKC412 treatment increased this ratio in all groups, but significance was reached only in control cells, and levels remained lower in *AXL*-expressing cells **(Figure 5G)**.

### Dual PKC412 and R428 treatment reduces the viability of *KIT* D816V/*AXL*-expressing MCs

We investigated dual targeting of KIT and AXL with PKC412 and R428 (bemcentinib), a selective but not fully specific AXL inhibitor. Cell viability was assessed by WST-8 assay over seven days, given the limited effect of PKC412 alone at 96 hours. Parental ROSA *KIT* D816V cells and counterparts expressing WT or L197M *AXL* were treated with 200 nM PKC412, 500 nM R428, the combination or DMSO. By day 7, PKC412 alone decreased viability by ∼80% in control cells but only by ∼50% in WT- or L197M *AXL-* expressing cells. R428 alone decreased viability by ∼40% in controls and by ∼20% and ∼10% in WT- and L197M *AXL-* expressing cells, respectively. Strikingly, the combination reduced viability by ∼70-80% in control and WT *AXL*-expressing cells and by ∼60% in L197M *AXL*-expressing cells **(Figure 6A)**. Thus, dual KIT and AXL targeting was more effective than either agent alone (see representative confluence images after seven days of treatment in **Figure 6B)**. Molecular analysis of apoptotic regulators revealed a modest decrease in BCL-XL expression with the combination treatment relative to PKC412 alone, whereas BCL2 and MCL1 remained unchanged in all conditions. The cleaved/procaspase-3 ratio tended to be higher with the combination in all groups **(Figure 6C)**. Survivin was unaffected (data not shown), but Livin and cIPA1 levels were lower with the combination than with PKC412 alone; this effect being less pronounced in *AXL* L19M-expressing cells, consistent with their partial resistance to dual treatment **(Figure 6D)**.

**Fig. 6.**
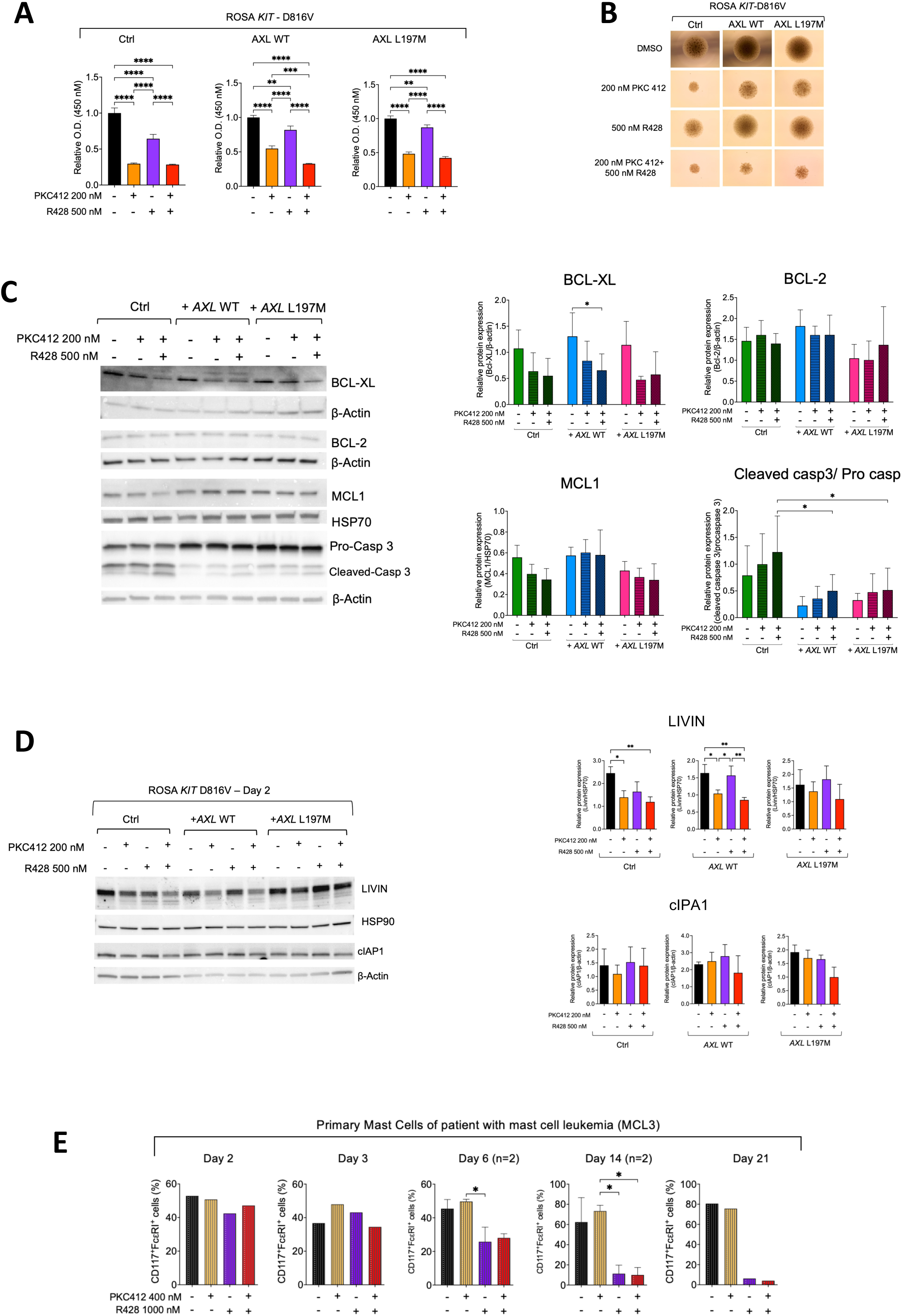
Effects of single and combination treatments on abnormal mast cells. ***(A)*** Cell viability of ROSA *KIT* D816V cells (control, *AXL* WT, or *AXL* L197M) measured by WST-8 assay after7 days of treatment with DMSO, 200 nM PKC412, 500 nM R428 or their combination. Viability was determined by absorbance at 450 nm. ***(B)*** Representative images of cell confluence at day7. ***(C)*** Immunoblots analysis of BCL2, BCL-XL, MCL1, Caspase 3 performed in cells after 48 h of treatments as in A. β-actin or HSP70 served for normalization. ***(D)*** Expression of Livin and cIPA1 assessed by immunoblots in cells treated for 48 hours, as indicated. Data are representative of three independent experiments. ***(E)*** Fresh PBMCs from patient MCL3 were treated with DMSO, 400 nM PKC412, 1 µM R428, or their combination. MCs viability (CD117⁺, FcεRI⁺, Sytox Blue⁻) was monitored by FACS over 21-days. Due to limited cell availability, only days 6 and 14 were analyzed in duplicate. Statistical significance was assessed by one-way ANOVA (A, D, E). Caspase analyses were performed by two-way ANOVA (C). *p < 0.05; **p < 0.01; ***p < 0.001; ****p < 0.0001.

### R428 alone triggers neoplastic MC death in an MCL patient

We had the rare opportunity to analyze a fresh peripheral blood sample from a mast cell leukemia (MCL3) patient with 27% circulating MCs carrying *KIT* F522C, *SF3B1* and *TERT* mutations. The patient was refractory to midostaurin (PKC412). FACS revealed strong AXL expression in all MCs **(Figure 2B)**. PBMCs were immediately treated with 400 nM PKC412, 1 μM R428, the combination, or DMSO. Limited cell number availability restricted duplicate testing to days 6 and 14. MC viability was monitored by FACS for three weeks, with medium and drug renewal every three days. Consistent with clinical data, PKC412 had no effect on MCs. By contrast, R428 monotherapy induced ∼50% cell death by day 6, ∼70% by day 14, and ∼90 % by day 21. The combination was no more effective than R428 alone **(Figure 6E)**. These findings highlight the therapeutic potential of AXL inhibition in AdvSM patients with non-D816V *KIT* mutations.

## Discussion

The unexpected expression of AXL in neoplastic MCs identifies this receptor as a novel player in mastocytosis pathophysiology. Given its established role in cancer, AXL emerges as a potential driver of mastocytosis progression and TKI resistance.

We detected AXL in neoplastic MCs from patients with various forms of mastocytosis. One patient carried a germline L197M mutation in the *AXL* gene. Expression levels varied across patients and tissues, whereas public scRNA-seq datasets showed that *AXL* is largely absent from MCs in healthy skin, BM, and non-mastocytosis tumors. Together, these findings suggest that *AXL* expression in MCs is a disease-specific feature of mastocytosis.

Paradoxically, AXL was undetectable *in vitro* in mast cell lines and CD34^+^-derived MCs under steady-state conditions. However, its expression was induced by several stimuli, with type I interferons (IFN-α and IFN-β) being the most potent, indicating that inflammatory signals regulate *AXL* expression in MCs. Consistently, IFN levels were high in patient plasma, reinforcing the link between inflammation and *AXL* induction. Recent studies have also reported increased inflammatory cytokines levels in plasma from systemic mastocytosis, along with altered distributions of activated monocytes and dendritic cells potentially facilitating their recruitment to MC-enriched tissues. In these inflammatory contexts, *AXL* upregulation has also been observed in dendritic cells. While these studies focused primarily on antigen-presenting cells, the authors hypothesized that neoplastic MCs might contribute to a self-sustaining inflammatory loop ^28–29^. Our findings support this hypothesis, suggesting that an inflammatory microenvironment may promote *AXL* expression in MCs *in vivo* through STAT1 and pSTAT1 upregulation, as reported in other cellular models ^12,30^. In various cell types, type I IFNs induce *AXL*, leading to a limitation of their production through a negative feedback loop to prevent excessive immune responses. We can speculate that *AXL* expression in neoplastic MCs may similarly shape the immune landscape by facilitating immune evasion, although this lies beyond the scope of the present study ^7^. We also suggest that GAS6-mediated autocrine and paracrine activation contributes to AXL expression, as MCs produce this ligand and are exposed to it in the bloodstream. Plasma GAS6 concentrations did not differ between patients and healthy individuals, but local overexpression in mastocytosis lesions cannot be excluded. Hypoxia, a common feature of the hematopoietic niche and a known inducer of *AXL* in cancer ^7,31,32^, had no effect on MCs *in vitro,* leaving its role in mastocytosis uncertain.

Functionally, our findings support an oncogenic role for AXL in MCs. Both WT and L197M *AXL* enhanced proliferation and clonogenicity, but this effect was strictly dependent on *KIT* D816V, indicating that AXL amplifies KIT-driven oncogenic signaling. RNA-seq data from ROSA *KIT* D816V cells and their AXL-expressing counterparts confirmed these results, showing significant enrichment of pathways related to proliferation, viability and survival, with concomitant downregulation of cell death pathways.

This prosurvival effect was confirmed in IL-3–dependent BaF3 cells, in which both WT and L197M *AXL* promoted survival in the absence of IL-3. In this model, the L197M variant was more protective than WT, suggesting enhanced activity under stress conditions. Unlike ROSA *KIT* D816V cells, neither WT nor L197M *AXL* increased BaF3 cell proliferation. These results indicate that AXL exerts dual functions: promoting survival independently of other factors and enhancing proliferation exclusively in cooperation with oncogenic signals, such as *KIT* D816V. RNA-seq data from ROSA *KIT* D816V and *AXL*-expressing cells were consistent with these findings. Moreover, IPA on scRNA-seq datasets from normal tissues revealed an enrichment in survival- and viability-related pathways and a repression of apoptotic and necrotic signatures in *AXL*-expressing MCs. AXL expression may therefore confer a survival advantage even in normal MCs, if sustained **(Supplementary Figure 2)**. Mechanistically, *AXL* expression was associated with increased activation of STAT5, STAT3, p38, and FAK, together with upregulation of survivin, BCL2, MCL1, and BCL-XL, consistent with roles in proliferation, multifocal expansion, survival, and tumor progression.

*AXL* expression conferred resistance to PKC412, with sustained BCL2, MCL1 and BCL-XL expression and reduced caspase-3 activation, consistent with reports in AML and endothelial cells ^8,32^. Dual targeting of KIT D816V and AXL with PKC412 and R428 enhanced drug sensitivity, with a slightly stronger effect in WT-than in L197M *AXL*-expressing cells. This subtle difference suggests that the variant may confer a greater survival advantage *in vivo*. The dual treatment effect was associated with reduced BCL-XL levels and downregulation of LIVIN and cIAP1, which act downstream of caspase-3. Persistent BCL2 and MCL1 expression likely account for the residual viability.

Remarkably, we demonstrated high levels of *AXL* expression in neoplastic MCs from an MCL patient harboring *KIT F522C*, *SF3B1* and *TERT* mutations, who failed to respond to PKC412. Consistently, these MCs proved completely refractory to PKC412 *in vitro* but were strikingly sensitive to R428, with no additional benefit from its combination with PKC412. Notably, AXL expression persisted for more than two weeks in culture, suggesting regulation by autocrine/paracrine stimulation or by genetic/epigenetic mechanisms driving AXL overexpression.

**In conclusion,** our findings identify AXL as a key survival factor and major driver of TKI resistance in MCs, highlighting a critical therapeutic vulnerability in AdvSM. Dual KIT/AXL inhibition outperformed PKC412 alone in *KIT*-mutated cells. We also show, for the first time, that targeting AXL induces marked cytotoxicity in neoplastic MCs from a patient lacking *KIT* D816V, the major target of PKC412 and avapritinib, thereby positioning AXL as a therapeutic target in *KIT* D816V–independent mastocytosis. Nevertheless, despite these promising results, the persistence of BCL2 and MCL1 supports the need for additional strategies, and combined inhibition of KIT, AXL, and BCL2 family proteins may provide a path toward complete eradication of neoplastic MCs cells in patients.

## Abbreviations

AML: acute myeloid leukemia
AdvSM: advanced systemic mastocytosis
ASM: aggressive systemic mastocytosis
BM: bone marrow
BMMC: bone marrow mast cell
CML: chronic myeloid leukemia
CM: cutaneous mastocytosis
EMT: epithelial-to-mesenchymal transition
FDA: Food and Drug Administration
IFNα: interferon alpha
IFNβ: interferon beta
IFNγ: interferon gamma
IHC: immunohistochemistry
ISM: indolent systemic mastocytosis
TKI: tyrosine kinase inhibitor
IPA: Ingenuity pathway analysis
MC, MCs: mast cell, mast cells
MCL: mast cell leukemia
SCF: stem cell factor
WBM: whole bone marrow
WT: wild type

## Acknowledgments

We thank Sylvie Fabrega for assistance with virus production and Lucile Marchal for bioinformatics advice. We thank Cédric Broussard from the Proteom’IC platform at Institut Cochin for his contribution to the FAK analysis. We also thank Prof. François Moreau Gaudry for providing the pRRSLIN-MND-PGK-Tomato-WPRE backbone. This research was supported by grants from the Association Laurette Fugain (ALF), The French Society of Dermatology (SFD), the CEREMAST and INSERM Transfert. Kitsada Kangboonruang was funded by IDEX-Université Paris Cité-ED561, Imagine Institute and the CARNOT Imagine.

## Author contributions

L.M-C initiated the project and designed the study with the valuable contribution of O.H. Both analyzed and discussed the results. LM-C wrote the manuscript and OH provided insightful revision. KK conducted experiments, analyzed and discussed the results, generated the figures, wrote the Material and Methods sections and reviewed the manuscript. Y.L. and F.B. critically read the manuscript and provided comments. PD, TM performed IHC. FM, CK, SL, MF, GL, HH, MD participated in the experiments. MH assisted with colony formation assays. NC analyzed bioinformatics data. JM sorted the cells. ES, ND performed RNAseq. CB, CM, LP, TM, SB, CL, CM, LL, FR, DG, AL, MA, JR, FB, AB provided critical biological material. All authors approved the manuscript.

## Disclosure of Conflict of Interest

The authors declare no conflicts of interest for this study.

---

**Supp Fig 1:**
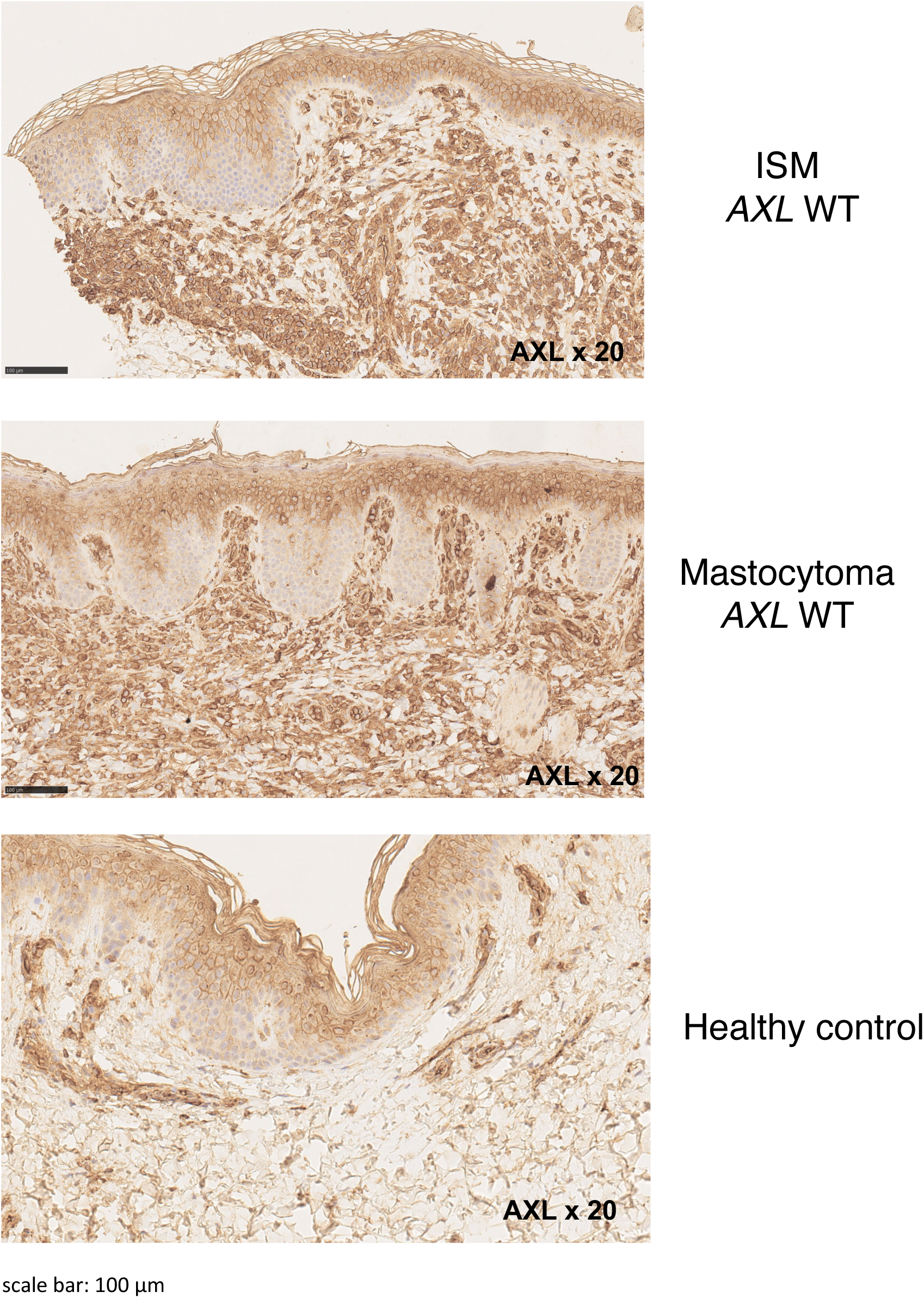
Histological analysis of skin sections from one ISM and one mastocytoma showing strong expression of AXL. Original magnification, ×20 (insets).

**Supp Fig 2:**
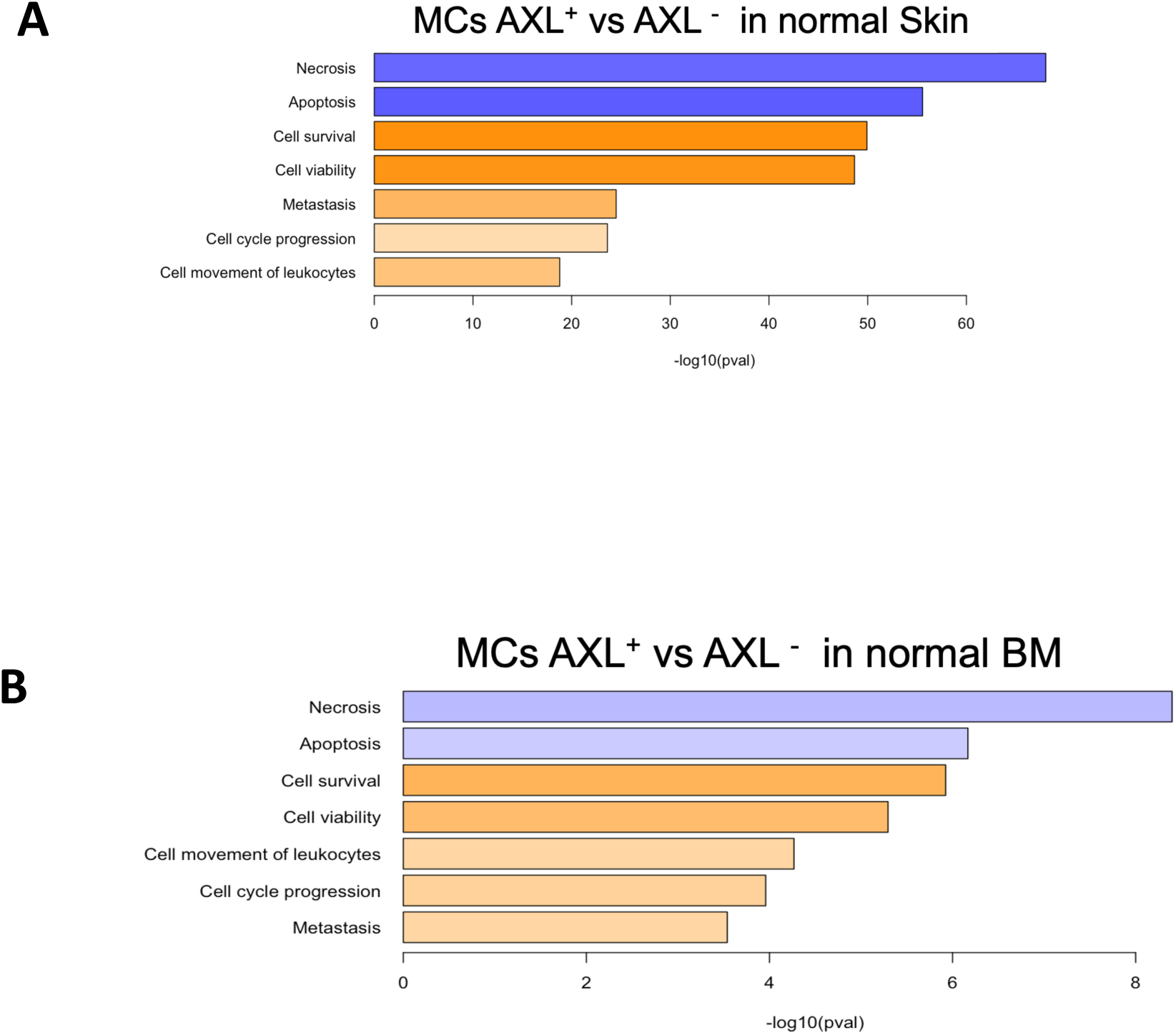
Ingenuity pathway analysis of AXL-positive versus AXL-negative MCs from normal Skin (A), normal BM (B). Blue= Decrease. Orange/pink= increase

## Supplemental methods

### Patient Samples

This retrospective study was conducted within the CEREMAST network. Bone marrow and skin samples (paraffin-embedded or frozen) were collected at diagnosis from patients classified according to the 2016 and 2022 WHO criteria and previously enrolled in an AFIRMM-supported prospective study. A fresh blood sample from one mast cell leukemia patient (MCL3) was processed for the isolation of PBMCs after ACK-mediated red blood cell lysis for downstream analyses. AXL expression was evaluated on all samples. The study was approved by the local ethics committee (*Comité de Protection des Personnes Ile-de-France*; approval 93-00) and conducted in accordance with the Declaration of Helsinki. All patients provided written informed consent. Patient details, diagnoses, and *KIT/AXL* status are summarized in **Table 1**.

### Immunohistochemical analysis of skin and BM biopsy specimens

Paraffin-embedded skin and BM sections were stained with H&E or subjected to IHC using anti-CD117 to assess MC infiltrates, and anti-AXL (AF154) (R&D Systems). Staining was performed with the Menarini refine detection kit

### Flow cytometric analysis of AXL expression in MCs

Ficoll-treated bone marrow (BM) samples from patients with mastocytosis were thawed from liquid nitrogen. Cells were collected and washed twice with phosphate-buffered saline (PBS) containing 0.5% bovine serum albumin (BSA) and 2 mM ethylenediaminetetraacetic acid (EDTA). After the final wash, the supernatant was discarded, leaving approximately 100 µL of residual volume, and the cell pellet was thoroughly dissociated by vortexing. Cells were fixed and permeabilized using the Foxp3/Transcription Factor Staining Buffer Set (eBioscience). Briefly, 1 mL of Foxp3 Fixation/Permeabilization working solution was added to each sample, followed by vortexing and incubation for 30 minutes at 2–8 °C in the dark. Samples were then washed twice with 2 mL of 1× Permeabilization Buffer (1300 rpm, 5 minutes, room temperature). The cell pellet was blocked with Fc-blocker (1:500 dilution) and subsequently stained with PE-conjugated anti-AXL (Invitrogen), APC-conjugated anti-CD117 (Invitrogen), and PE-Cy7–conjugated anti-FcεRI (Invitrogen) for 1 hour at 2– 8 °C in the dark. After staining, cells were washed twice with 1× Permeabilization Buffer. Finally, cell pellets were resuspended in PBS containing 0.5% BSA and 2 mM EDTA, and AXL expression in mast cells was assessed using a BD FACSAria™ II Flow Cytometer (BD Biosciences).

### Ultra-sensitive digital ELISA (Simoa) assays

IFN-α, IFN-β, and IFN-γ levels were quantified using Simoa digital ELISA (Quanterix Homebrew). Plasma samples were thawed, centrifuged at +4°C to remove debris and analysed. The IFN-α assay detects all subtypes with similar sensitivity. Detection limits were 0.6 fg/mL (IFN-α), 1.54 pg/mL (IFN-β), and 1.0 fg/mL (IFN-γ). Results were compared to healthy controls.

### Plasma GAS6 measurement

A Luminex assay was performed on plasma samples from 10 healthy individuals and 37 SM patients to measure GAS6 levels.

### RNA extraction and real-time PCR

Total RNA was extracted from human MC lines or CD34+-derived primary MCs, treated or not with IFN-α (100/300 ng/mL), IFN-β (1–25 ng/mL), or IFN-γ (10–400 ng/mL) using the RNeasy Mini Kit (Qiagen). The quality and quantity of RNA was determined using a NanoDrop 2000C (Thermo Scientific). Complementary DNAs (cDNAs) were synthesized by SuperScript^TM^ IV VILO^TM^ Master Mix (Invitrogen). Following reverse transcription, the expression of *AXL* was examined by real-time PCR using TaqMan Real-Time PCR Assays (LifeTechnologies) with the CFX96TM Real-Time System (Bio-Rad), normalizing mRNA levels to 18S rRNA. Experiments were run in triplicate, with data presented as mean ± SD. Statistical significance was assessed by one-way ANOVA. **Table 2** of TaqMan assays is provided in Supplementary Data.

### Mast cell lines

Two ROSA MC lines^16^, originally derived from CD34^+^ umbilical cord blood (ROSA-*KIT* WT and ROSA-*KIT* D816V), or HMC1.2 cells (*KIT* V560G/D816V) from an MCL patient were used in this study. BMMCs were obtained from *KIT* D814V transgenic mice. Only ROSA-*KIT* WT cells were SCF-dependent. All human MC lines were cultured in Iscove’s Modified Dulbecco’s Medium (Invitrogen), supplement with 10% fetal bovine serum (Gibco). BMMC-*KIT*-D814V were cultured in Opti-MEM, supplement with 10% fetal bovine serum (Gibco). All cells were maintained in a humidified 95% air incubator with 5% CO_2_ at 37 C°. All MC lines had their medium changed twice per week and were regularly tested for mycoplasma contamination using MycoStrip^TM^ (InvivoGen).

### Primary MCs

Primary MCs were derived from peripheral blood CD34^+^ cells cultured in Iscove’s Modified Dulbecco’s Medium, supplement with 15% BIT (contains BSA and Insuline /transferrin/sodium selenite) in the presence of human SCF (100 ng/ml), IL6 (50 ng/ml), and for the first week, IL3 (10 ng/ml). Primary MCs underwent a medium change once per week and differentiation was assessed by FACS using APC-conjugated anti-CD117 (Invitrogen), and PE-Cy7-conjugated anti-FcεRI (Invitrogen).

### Lentiviral vector production and cell transduction

Lentiviral vectors pRRSLIN-*AXL* WT-*tdTomato* and pRRSLIN-*AXL* L197M-*tdTomato* were generated using the pRRSLIN-MND-PGK-*tdTomato*-WPRE backbone (gift from Prof. François Moreau-Gaudry, France). ROSA *KIT* WT, ROSA *KIT* D816V, BMMC *KIT* D814V and primary MCs were transduced with lentiviral particles in the presence of LentiBOOST (Sirion Biotech) to enhance transduction efficiency and incubated overnight at 37°C in a humidified 5% CO₂ incubator. The following day, the medium was replaced with fresh complete growth medium. Positive cells were sorted based on tdTomato expression using BD FACSAria^TM^ II cell sorter (BD Biosciences).

### *In vitro* cell proliferation measurement

Cell proliferation kinetics of MC lines (ROSA *KIT* WT, ROSA *KIT* D816V, and BMMC *KIT* D814V) expressing *AXL* WT-*tdTomato*, *AXL* L197M-*tdTomato* or control, were monitored in real-time using the Incucyte® live-cell imaging system (Sartorius) according to the manufacturer’s protocol. Briefly, cells were seeded in 96-well plates coated with Poly-L-ornithine solution at a density of 10,000 cells per well in complete growth medium. Plates were placed inside the Incucyte system maintained at 37°C in a humidified atmosphere containing 5% CO₂. Phase-contrast images were captured every 12 hours using a 10× objective lens. Cell proliferation was quantified by measuring the percentage of confluence over time using the Incucyte® integrated software. All data are representative of 3 independent experiments. Statistical significance was assessed by one-way ANOVA.

### *In vitro* cell proliferation assay on cells treated with a TK inhibitor

ROSA *KIT* WT and ROSA *KIT* D816V cells expressing AXL *WT*–tdTomato, *AXL* L197M–tdTomato, or control were treated with DMSO (vehicle) or 200 nM PKC412. Cell confluence was monitored for seven days using the IncuCyte system.

### Colony formation assay

ROSA *KIT* D816V cells expressing *AXL* WT-*tdTomato*, *AXL* L197M-*tdTomato*, or Ctrl (40,000 cells) were plated in a two-layer soft agar system (0.8% bottom, 0.48% top) using UltraPure^TM^ Low Melting Point Agarose (Invitrogen) and incubated at 37°C for over three weeks with weekly medium renewal. Colony formation was analyzed using ImageJ.

### Ba/F3 cell viability *in vitro*

Murine IL-3-dependent BaF3 pro-B cells, which lack endogenous *AXL* and *KIT* expression, were transduced with lentiviral vectors expressing *AXL* WT, *AXL* L197M or an empty vector control and cultured in the presence or absence of IL3. Cells were used to seed plates at a density of 20,000 cells/well and were cultured in RPMI-1640 medium with or without IL-3 (10 ng/mL). After 48 hours, cell viability was assessed in the Alamar Blue assay, according to the manufacturer’s instructions. The results presented are the mean ± SD of three independent experiments.

### RNA-seq Library Preparation from Microdissected Cutaneous Mast Cells

MCs from the skin lesions of 15 mastocytosis patients (12 ISM, 3 AdvSM) were microdissected, and RNA was extracted. The RNA integrity (RNA Integrity Score ≥ 7.0) was checked on the Agilent Fragment Analyzer (Agilent Technologies) and quantity was determined using Nanodrop. SureSelect Automated Strand Specific RNA Library Preparation Kit was used according to manufacturer’s instructions with the Bravo Platform (Agilent Technologies). Briefly, 100 ng of total RNA per sample was used for poly-A mRNA selection using oligo(dT) beads and subjected to thermal mRNA fragmentation. The fragmented mRNA samples were subjected to DNA synthesis and were further converted into double stranded DNA using the reagents supplied in the kit, and the resulting dDNA was used for library preparation. The final libraries were indexed, purified, pooled together in equal concentrations and subjected to paired-end sequencing (2×100 bp) on Novaseq-6000 sequencer (Illumina) at Gustave Roussy.

### In vitro cell death analysis

Cell death was assessed by FACS with SYTOX® Blue staining (Invitrogen) in ROSA *KIT* D816V cells expressing AXL *WT*–tdTomato, *AXL* L197M–tdTomato, or control treated with 200 nM PKC412 (200 nM) or DMSO (vehicle) for 96 hours. Cells were collected and stained with 1 uM of Sytox blue stain for at least 5 minutes at room temperature, protected from light. Analyze sample without washing or fixing by FACS with 440/40 nm bandpass filter.

### WST-8 Cell Viability Assay

ROSA *KIT* D816V cells (control, *AXL* WT, or *AXL* L197M) were seeded in 96-well plates at a density of 10,000 cells per well and treated for 7 days with DMSO, 200 nM PKC412, 500 nM R428, or a combination of both inhibitors. Cells were cultured at 37°C in a humidified atmosphere containing 5% CO₂. Cell viability was assessed using the WST-8 assay (Dojindo). Briefly, 10 ul of WST-8 solution was added to each well, followed by incubation for 4 hours at 37°C in the dark. Absorbance was measured at 450 nm using a microplate reader.

### Immunoblotting analysis

Cells were harvested and lysed in RIPA buffer (50 mM Tris-HCl pH7.4, 150 mM NaCl, 1.0% (v/v) NP-40, 0.5% (w/v) Sodium Deoxycholate, 0.1% (w/v) SDS, and 0.01% (w/v) sodium azide at a pH of 7.4) supplemented with and protease and phosphatase inhibitor cocktail. Lysates were incubated on ice for 45 minutes with intermittent vortexing, followed by centrifugation at 16,000 × g for 10 minutes at 4°C. The supernatants were collected, and protein concentration was determined using Pierce^TM^ BCA Protein Assay Kits (Thermo Scietific) according to the manufacturer’s instructions. Equal amounts of proteins (50 µg) were mixed with 4× Laemmli buffer and heated at 95 C° for 5 minutes. Proteins were separated on 4-15% sodium dodecyl sulfate polyacrylamide gel electrophoresis (SDS-PAGE) gels at constant voltage of 100-120 V in running buffer until the front dye reached the bottom of the gel. Proteins were subsequently transferred from the gel to a PVDF membrane (previously activated in absolute ethanol) by electro-blotting. Non-specific binding was blocked by incubating the membrane with 5% non-fat dry milk or 5% BSA (for phospho-proteins) in tris-buffered saline, 0.1% tween (TBST) for 1 hour at room temperature with gentle agitation. Membranes were incubated with the indicated primary antibodies diluted in blocking buffer overnight at 4 C° with gentle agitation and then washed 3 times for 10 minutes with TBST at room temperature with shaking. Membranes were incubated with HRP-conjugated secondary antibody (1:10,000 - 1:50,000 dilution in blocking buffer) for 1 hour at room temperature with gentle agitation. After washing with TBST 3 times for 10 minutes, protein bands were visualized using enhanced chemiluminescence (ECL) substrate according to the manufacturer’s instructions. The chemiluminescence signals were detected using ChemiDoc Imaging Systems (Bio-Rad). The densitometry analysis was performed using ImageJ software (NIH) and normalized to corresponding loading controls such as β-actin or HSP70.

Immunoblotting was performed on protein extracts from ROSA and HMC1.2 cells, with or without IFN stimulation, using anti-AXL (8661, Cell Signaling). ROSA *KIT* D816V cells expressing *AXL* WT-tdTomato, *AXL* L197M-tdTomato, or control were analyzed for the expression of STAT3 (9139, Cell Signaling), pSTAT3 (Y705) (9145, Cell Signaling), STAT5 (25656, Cell Signaling), pSTAT5 (Y694) (9359, Cell Signaling), FAK (3285, Cell Signaling), pFAK (Y397) (8556, Cell Signaling), p38α (sc-81621, Santa cruz), p-p38 (sc-7973, Santa cruz). Following 48 hours of treatment with PKC412 (200 nM), or DMSO, expression levels of BCL-XL (sc-56021, Santa cruz), BCL-2 (M0887, Dako), MCL1 (ab32087, Abcam), and caspase-3 (sc-7272, Santa cruz) were assessed. Protein loading was normalized using β-actin (sc-47778, Santa cruz).

### MCs from an MCL3 Patient treated with TKIs

Fresh PBMCs from the MCL3 patient were cultured in the presence of SCF and IL6 to selectively promote mast cell culture. One fraction was immediately treated with 200 nM PKC412, 500 nM R428, the combination of both, or DMSO (diluent) for 3 weeks. Media and treatments were refreshed every 3 days. MCs survival was assessed by flow cytometry using anti-CD117, anti-FcεRI, and Sytox Blue staining. Due to limited cells, not all treatment points were duplicated.

### Transcriptomic analyses

RNA-seq was performed on RNA extracted from parental ROSA KIT D816V cells and counterparts expressing AXL WT or AXL L197M. RNA-seq libraries were prepared using the Universal Plus mRNA-Seq kit (Nugen). cDNAs were sequenced on a NovaSeq6000 from Illumina. RNA-seq data generated during the current study are available in BioStudies and are publicly available at : https://www.ebi.ac.uk/biostudies/studies/S-BSST2177. FASTQ files were mapped to the Human GRCh38/hg38 reference using star and counted by featureCounts from the Subread R package. Duplicate reads were marked and removed during the counting process. Read count normalisations and groups comparisons were performed by three independent and complementary statistical methods : DESeq2 (v1.24.0), edgeR (v3.26.8), Limma Voom (v3.40.6). Flags were computed from counts normalized to the mean coverage. All normalized counts <20 were considered as background (flag 0) and >=20 as signal (flag=1). P50 lists used for the statistical analysis regroup the genes showing flag=1 for at least half of the compared samples. The results of the three methods were filtered at pvalue<=0.05 and folds 1.2/1.5/2 compared and grouped by Venn diagram. Hierarchical clusterings were performed using the Spearman correlation similarity measure and ward linkage algorithm. Functional analyses were carried out using Gene Ontology (org.Hs.eg.db v3.8.2) and Ingenuity Pathway Analysis (IPA, Qiagen). R (v3.6.1) from the R project for Statistical Computing [http://www.r-project.org/]

Ingenuity Pathways: [http://www.ingenuity.com]

### Statistical analysis

Quantitative data were presented as means ± SD. The statistical analysis was analyzed by using the statistical software package, GraphPad Prism 10 (GraphPad Software, San Diego, CA, USA). Statistical significance was determined by using unpaired 2-sample t tests or one-way ANOVA or two-way ANOVA (equal variance), *p*-Values <0.05 were considered as statistically significant. All data were normally distributed, and variance was similar in groups that were compared in statistical tests.

## Notes

https://www.ebi.ac.uk/biostudies/studies/S-BSST2177

### Competing Interest Statement

The authors have declared no competing interest.

